# Adrenomedullin-RAMP2 enhances endothelial cell homeostasis synergically with shear stress

**DOI:** 10.1101/2025.09.21.677608

**Authors:** Yongdae Yoon, Sean Duffy, Shannon Kirk, Kamoltip Promnares, Pratap Karki, Anna A Birukova, Konstantin G Birukov, Yuan Yuan

**Affiliations:** Department of Anesthesiology, University of Maryland School of Medicine, Baltimore, MD; Section of Pulmonary, Critical Care and Sleep Medicine, Department of Medicine, Yale School of Medicine, New Haven, CT; Pulmonary and Critical Care Medicine, University of Maryland School of Medicine, Baltimore, MD

## Abstract

Analysis of pulmonary vascular dysfunction in various lung pathologies remains challenging due to the lack of functional *ex vivo* models. Paracrine signaling in the lung plays a critical role in regulating endothelial maturation and vascular homeostasis. Previously, we employed single-cell RNA-sequencing (scRNAseq) to systematically map ligand-receptor (L/R) interactions within the lung vascular niche. However, the functional impact of these ligands on endothelial biology remained unknown. Here, we systematically evaluated selected ligands in vitro to assess their effects on endothelial barrier integrity, anti-inflammatory responses, and phenotypic maturation. Among the top soluble ligands, we found that adrenomedulin (ADM) exhibited superior barrier enhancing effect on human pulmonary endothelial cell monolayers, as evidenced by electrical cell impedance sensing (ECIS) and XperT assays. ADM also exhibited anti-inflammatory properties, decreasing ICAM1 and increasing IkBa expression in a dose-dependent manner. Shear stress (15 dynes/cm^2^) alone increased endothelial characteristics, including homeostatic markers such as *CDH5*, *NOS3*, *TEK*, and *S1PR1*. ADM treatment maintained the enhanced level of these markers under shear stress and further improved anti-coagulation by increasing *THBD* and decreasing *F3* expression, and synergistically enhanced the expression of the native lung aerocyte capillary endothelial marker *EDNRB*. This effect was completely attenuated by a blockade of ADM receptor, RAMP2. Together, these findings identify ADM/RAMP2 signaling as a key paracrine pathway that enhances vascular barrier integrity, anti-inflammatory phenotype, and endothelial homeostasis, providing a framework for improving the physiological relevance of engineered vascular models.

## 1. Introduction

Pulmonary vascular disease is a growing global health concern and a significant cause of morbidity and mortality ^1^. This condition is characterized by microvascular remodeling and rarefaction and is primarily attributed to pathological changes in the vascular endothelium ^2^.

Current in vitro systems, including organ-on-chips and organoid platforms, have advanced our ability to model vascular systems by incorporating differentiated or mature endothelial cells into hydrogel or polydimethylsiloxane (PDMS)-based perfused systems. These models replicate aspects of physiological flow and intercellular communication ^3–6^, providing valuable insights into disease mechanisms. However, these platforms fall short of replicating the complex physiology of the human lung vasculature and lack critical microenvironmental cues in the vascular niche that is essential to alveolar endothelial function, including paracrine signaling, matrix substrate components, and correct substrate stiffness. As a result, no in vitro system can recapitulate native pulmonary vascular cellular phenotypes and physiological functions over extended periods in a controlled manner.

During normal physiological conditions, local cell-cell interactions among endothelial, epithelial, stromal, and immune cell populations within the lung vascular niche can jointly facilitate endothelial homeostasis and protect vasculature from injury. For example, cell-cell crosstalk paracrine factors, including angiopoietin-1 (Ang-1), and slit guidance ligand 2 (Slit2), are secreted in the vascular niche to enhance barrier function and reduce inflammation, preventing the influx of cells and plasma from the bloodstream into the local tissues (reviewed in ^7^). To investigate these crosstalk signals under homeostatic conditions, we previously employed single-cell RNA-sequencing (scRNA-seq) to identify cellular populations in human lungs and to characterize signaling molecules secreted by alveolar epithelial, mesenchymal, and immune cells that may be sensed by endothelial cells ^8^. We mapped putative ligand-receptor (L/R) interactions within and between cell types and assessed the significance of these interactions using the NicheNet database ^8^. From this analysis, we identified a subset of L/R pairs, including *ADM/RAMP2*, *SLIT2/ROBO4*, *BMP5/BMPR2*, that are highly enriched in the lung vascular niche and potentially interact with receptors expressed by vascular endothelial cells and contribute to vascular homeostasis. In this study, we assessed the impact of these top ligands on endothelial cells and tested their ability in regulating vascular cellular phenotypes and physiological functions in the in vitro vascular model systems.

The endothelium lining the blood vessels is highly sensitive to hemodynamic forces that act at the vascular luminal surface in the direction of blood flow (reviewed in ^9^). It was proposed that endothelial cells have their preferred fluid shear stress, or ‘set point’ ^10^. Upon exposure to 16 hours of laminar shear stress ranging from 2 to 60 dynes/cm^2^, human umbilical vein endothelial cells (HUVECs) aligned in the direction of the flow and anti-inflammatory and endothelial quiescence pathways were activated in the range of 10 to 20 dynes/cm^2^, but cells were misaligned or oriented perpendicularly (against the flow direction), and inflammation and coagulation signaling was activated outside this range (^10^ and reviewed in ^11^). In this study we investigated the impact of the top ligands involved in paracrine signaling in the intact lung, on cell-cell junction, anti-inflammatory response, anti-coagulation, and native cellular phenotypes in both static and dynamic flow conditions at a shear stress of 15 dynes/cm^2^ in order to identify key players preserving pulmonary endothelial homeostasis that may improve the fidelity of the engineered in vitro models of the pulmonary vasculature.

## 2. Methods

### 2.1. Materials

Adrenomedullin (ADM, #24889), Secretin (24990), and Transforming growth factor beta1 (TGF-β1) from Cayman Chemical (MI, USA); Slit homolog 2 (SLIT2, # 8616-SL), Semaphorin 6D (Sema6D, 2095-S6), Pleiotrophin (252-PL), vascular endothelial growth factor A (VEGFA, BT-VEGF), C-C motif chemokine ligand-19 (CCL19, 361-MI/CF), CXC Motif Chemokine Ligand-6 (CXCL6, 333-GC/CF), and TNF-α (# 10291-TA) from R&D Systems (MN, USA); Angiopoietin-1 (ANGPT1, #923-AN) and Bone morphogenic protein 5 (BMP5, # 615-BMC-020) and BMP3 (NBP2-61322) from Novus Biologicals (CO, USA); and BMP7 (595602), BMP6 (595502), BMP4 (59202), and BMP-2 (767302) were purchased.

### 2.2. Cell culture

Human pulmonary artery endothelial cells (HPAECs) were purchased from Lonza (Basel, Switzerland). The cells between passage 6-7 were placed on fibronectin 1 µg/cm^2^ coated plates and were incubated in EGM-2 endothelial cell growth medium-2 bulletkit from Lonza under a cell culture incubator at 37 with 5% CO_2_ and a humidified atmosphere.

### 2.3. Electrical cell-substrate impedance sensing

Electrical cell-substrate impedance sensing (ECIS) assays were performed according to the manufacturer’s instructions. Briefly, ECIS Cultureware plates (e.g., 96W20idf for static conditions or 1F8x10E chips for shear stress) from Ibidi (Grafelfing, Germany) were incubated with L-cysteine (10 mM) and then coated with fibronectin (1 µg/cm^2^). For static conditions, cells were incubated for 2 days until a stable baseline impedance was achieved. Ligands were then added to the cells, and changes in impedance were measured to determine endothelial cell permeability. For shear stress conditions, cells were incubated for 1 day after attachment. An unidirectional shear stress of 15 dynes/cm^2^ was applied to the cells, and ADM was added for one more day with continuous impedance measurement.

### 2.4. Expression permeability test assay

To verify endothelial cell permeability to macromolecules, expression permeability test (XperT) assay was performed, as previously described^12^. In brief, cells were seeded on plates coated with biotinylated collagen type I (0.25 mg/mL) dissolved in a 0.1 M bicarbonate, pH 8.3 solution. After ligand treatment, the cells were incubated with FITC-avidin (25 µg/mL) for 2 minutes. Unbound FITC-avidin was removed by washing with PBS. The fluorescence signal was then either visualized using an EVOS microscope (Thermo Fisher Scientific, MA, USA) or measured with a SpectraMax® iD5 microplate reader (Molecular Devices, CA, USA).

### 2.5. Western blot

Cells were lysed with 2x Laemmli sample buffer (#1610737, Bio-Rad, CA, USA). The lysates were boiled for 5 minutes and then subjected to SDS-polyacrylamide gel electrophoresis (SDS-PAGE). Proteins were transferred onto a polyvinylidene difluoride (PVDF) membrane (Bio-Rad). The membrane was blocked with 5% skim milk or 5% BSA in Tris-HCl-buffered saline containing 0.05% Tween 20 (TBST) for 30 minutes. It was then incubated with primary antibodies against ICAM-1 (sc-8439) and GAPDH (sc-47724) from Santa Cruz Biotechnology (Santa Cruz, CA, USA), as well as IκBα (#4814), NF-κB p65 (#3032), and phospho-NF-κB p65 Ser536 (#3031) from Cell Signaling Technology (MA, USA). Primary antibody incubation was performed overnight at 4°C. Following primary antibody incubation, the membrane was incubated with horseradish peroxidase (HRP)-conjugated secondary antibodies (7074S and 7076S, Cell Signaling Technology) for 2 hours. After washing the membrane three times with TBST, the protein bands were visualized using a ChemiDoc XRS+ System (Bio-Rad) after treatment with SuperSignal West Pico PLUS or Atto Ultimate sensitivity chemiluminescent substrate (34578 or A38556, Thermo Fisher Scientific).

### 2.6. qPCR

Total RNA was purified using the RNeasy Plus Mini Kit (#74136, Qiagen, Hilden, Germany) according to the manufacturer’s instructions. Complementary DNA (cDNA) was synthesized from the purified RNA using an iScript cDNA Synthesis Kit (#1708890, Bio-Rad). For real-time PCR, the cDNA was mixed with gene-specific primers and PerfeCta SYBR Green FastMix (Quantabio, MA, USA). The reactions were run on a CFX Opus 384 Real-Time PCR System (Bio-Rad). The primer sequences are listed in Table 1. Relative mRNA expression was quantified using the 2^-(△△Ct)^ method.

**Table 1.**
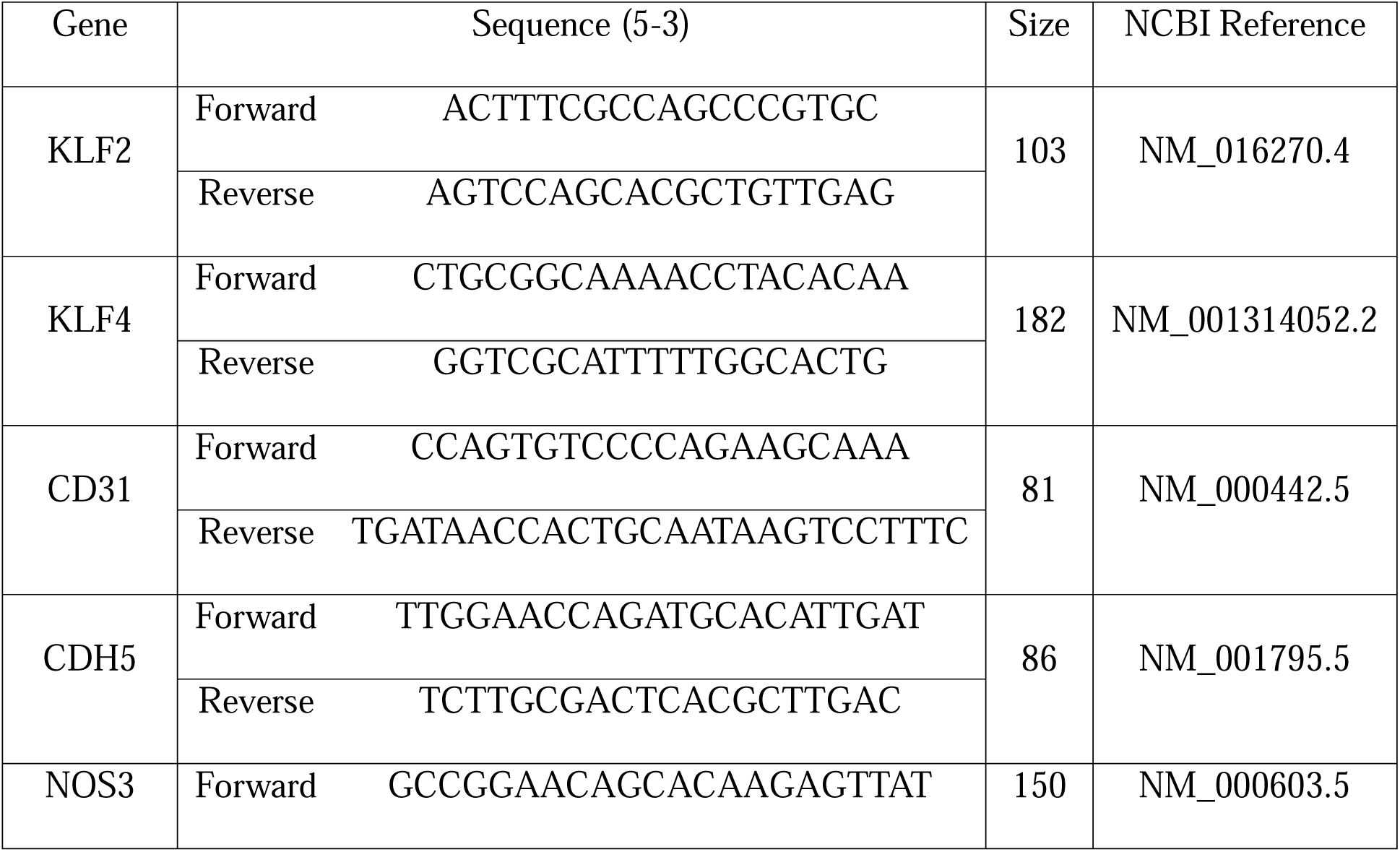
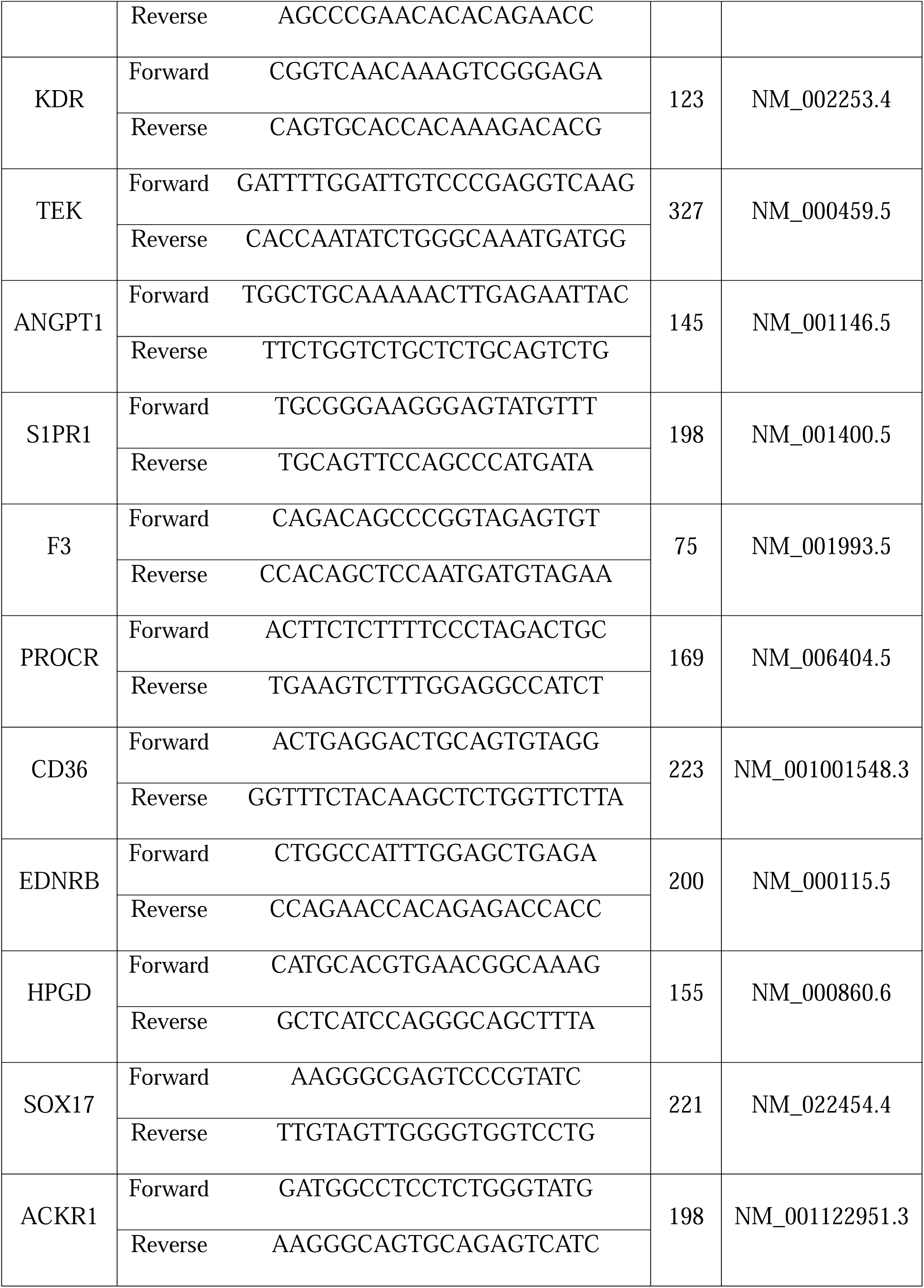
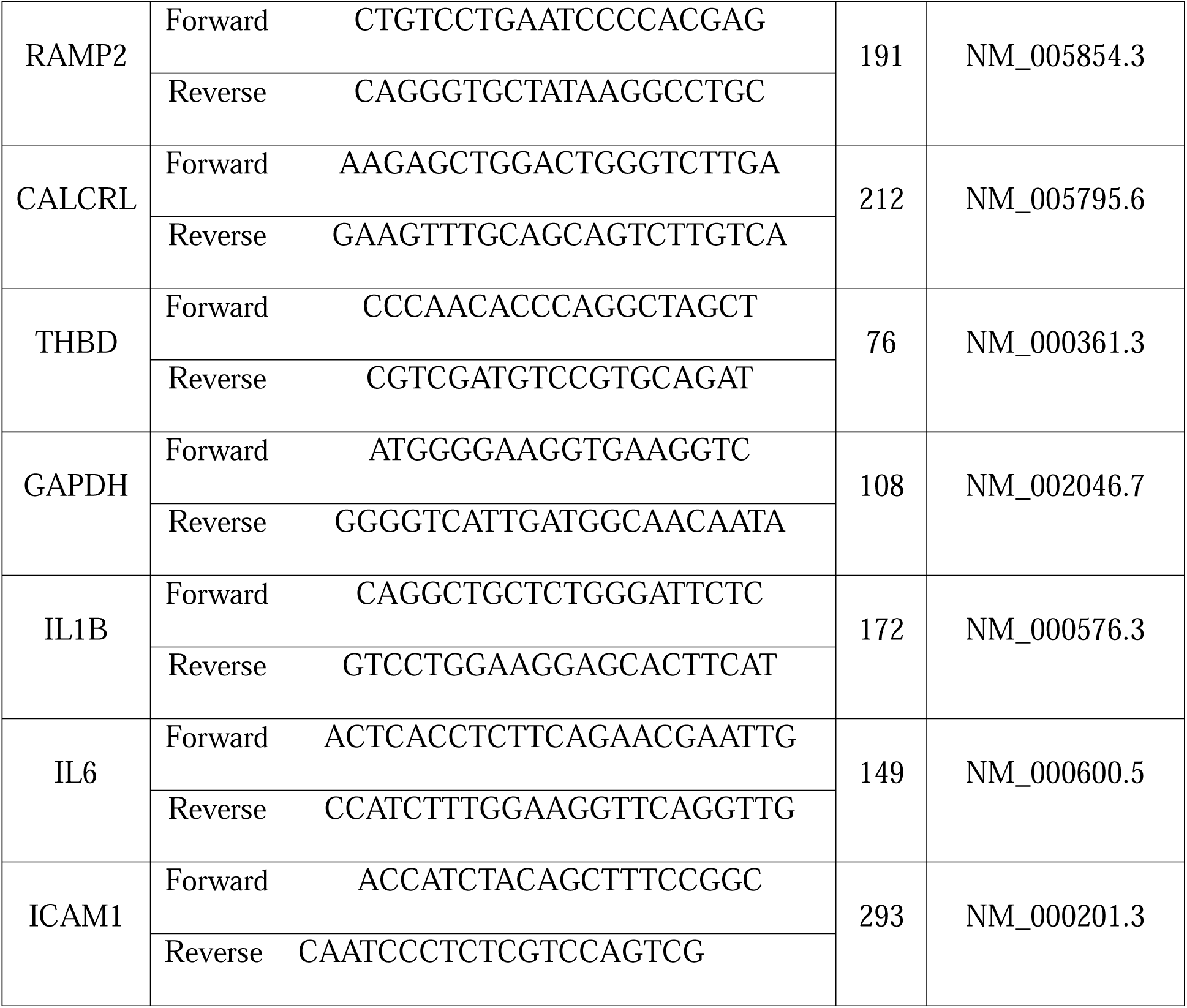
Primer list.

### 2.7. Enzyme-linked Immunosorbent Assay (ELISA)

The levels of soluble intercellular adhesion molecule-1 (ICAM-1) in the culture medium were measured using a human ICAM-1 ELISA kit (#DY720) and a Clear Microplate (#DY990) from R&D Systems (MN, USA). Cells were treated with TNF-α (1 ng/mL) for 30 minutes, followed by treatment with ADM (100 ng/mL) for 6 hours. The assay was performed according to the manufacturer’s instructions. The values were quantified by measuring absorbance at 450 nm using a SpectraMax® iD5 (Molecular Devices, CA, USA).

### 2.8. Shear Stress

To apply an unidirectional shear stress to endothelial cells, we utilized an ibidi pump system (#10902, ibidi, Gräfelfing, Germany). The cells were seeded on µ-Slide 0.4 Luer ibiTreat (#80176, Ibidi) coated with fibronectin (1–2 µg/cm^2^). Following the manufacturer’s instructions, the slides were connected to the pump system. The cells were subjected to 15 dynes/cm^2^ shear stress for 1 day. Subsequently, the medium was carefully replaced with fresh medium with or without ADM (100 ng/ml), and the cells were incubated for an additional day under shear stress at 15 dynes/cm^2^.

### 2.9. Immunofluorescence

For static condition, HPAECs were cultured on glass coverslips coated with fibronectin at 2 µg/cm^2^. For shear stress condition, cells were seeded in a µ-Slide ^0.4^ Luer ibiTreat chip (#80176, Ibidi) with fibronectin coating. Cells in both conditions were fixed with 2% paraformaldehyde in PBS including Ca^2+^ and Mg^2+^ and permeabilized with 0.1% Triton X-100 in PBS for 15 min at RT. Following a 30 min incubation with 5% BSA in PBS, cells were incubated with VE-cadherin antibody (#93467, Cell Signaling Technology) in 5% BSA in PBS at 4 °C overnight. The cells were then labeled by an Alexa Fluor^TM^ Plus 488-conjugated secondary antibody (Invitrogen, CA, USA) for 1h at RT. For filamentous actin staining, the cells were incubated with Texas Red-X phalloidin (#T7471, Thermo Fisher Scientific) for 30 min at RT. After DAPI staining (#62248, Thermo Fisher Scientific), the stained cells were mounted by a ProLong glass antifade Mountant. Images were captured using a fluorescent microscope (Eclipse TS2R, Nikon, Tokyo, Japan).

### 2.10. siRNA transfection

Transfection of RAMP2 (sc-36378) or control siRNA (sc-37007) from Santa Cruz Biotechnology (Santa Cruz, CA, USA) was performed using Lipofectamine RNAiMAX Transfection Reagent (13778-075, Thermo Fisher Scientific, MA, USA) according to the manufacturer’s instructions. Briefly, HPAECs were seeded in fibronectin-coated 6-well plates. Cells were then transfected with either RAMP2 or control siRNA. Following a 24-hour incubation, the transfected cells were replated onto ECIS plates for static conditions, ECIS chips for shear stress experiments, or µ-Slide Luer ibiTreat chips for immunofluorescence.

### 2.11. Statistical analysis

Data are presented as the mean ± standard deviation (SD) from three independent experiments. Statistical analysis was performed using GraphPad Prism 10 software. Differences between two groups were analyzed with an unpaired Student’s t-test. A p-value of less than 0.05 was considered statistically significant. *p<0.05; **p<0.01; and ***p<0.001

## 3. Results

### 3.1. ADM improves endothelial barrier function

To determine the impact of native soluble ligands expressed in human lung vascular niche on endothelial homeostasis, we leveraged our L/R pair database previously generated from scRNAseq data of human lungs ^8^. We focused our analysis on the top ligands, including ADM, Ang-1, Slit2, Secretin, Sema6D, Pleiotrophin, TGF-β1, VEGF-A, CCL19, CXCL6, BMP2, BMP5, BMP4, BMP6, and BMP7, expressed by alveolar epithelial, mesenchymal, and immune cells, within the vascular niche that could interact with receptors on endothelial cells identified through our previous *Connectome* analysis (Fig. 1A and ^8^). Among the ligands tested, we found that both ADM and Ang-1 enhanced endothelial cell barrier integrity, with ADM exhibiting the strongest effect in HPAECs. In contrast, VEGF-A strongly, and CXCL-6, CCL-19, and BMP5 modestly impaired the endothelial barrier (Fig. 1B), while all other ligands had no or negligible impact on cell-cell junction. Additionally, the XperT and ECIS assays consistently demonstrate ADM increased barrier function in a dose-dependent manner in HPAECs, especially at 10 and 100 ng/mL (Figs. 1C and D). These results demonstrate that the soluble ligands identified from native human lung niche might have a distinct impact on endothelial barrier function, and ADM exerts a superior barrier-protective impact as compared to all other tested ligands in vitro cell culture models.

**Figure 1.**
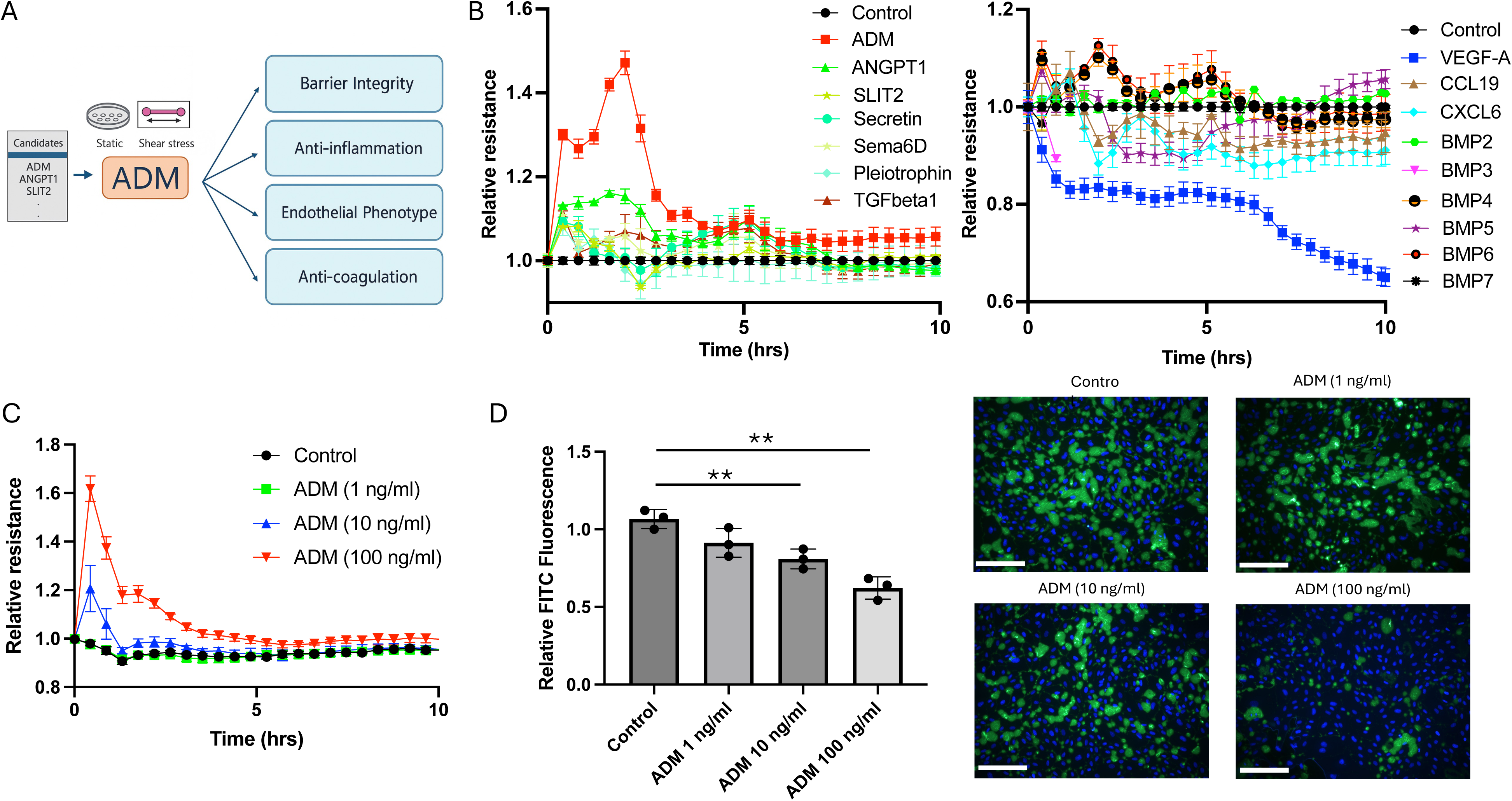
ADM exhibits superior effect on enhancing vascular endothelial barrier function. (A) The image presents an overview of experimental flow, including a list of candidate ligands that were selected from a preliminary analysis of ligand/receptor pair between endothelial cells and non-endothelial cells in macrovascular environments. (B) Barrier function of HPAEC monolayers was tested by ECIS assay after the cells were treated with the selected ligands. (C) Dose-dependent effect of ADM (1 - 100 ng/ml) on HPAECs was measured using ECIS assay. (D) HPAECs were seeded on biotinylated collagen type I (0.25 mg/mL) coated well plates. The cells were pretreated with the indicated ADM concentrations for 1 hour, and FITC-conjugated avidin was added for 2 minutes. The FITC fluorescence was measured, and the image of FITC signal was captured after DAPI staining. The graph is presented as the means ± standard deviation (SD) from three independent experiments. White scale bars: 150 μm, **\*\***p<0.01

### 3.2. ADM reduces inflammatory responses

To determine the impact of ADM on anti-inflammatory responses, HPAECs were pre-treated with TNF-α for 30 minutes prior to ADM treatment. We found that ADM significantly decreased the ICAM-1 expression and restored the degradation of IκBα, an inhibitory enzyme that reduces the NF-κB activation, after TNF-α challenge (Figs. 2A and B). Consistently, treatment with ADM reduced the phosphorylation of p65 at S536, a subunit of NF-κB (Fig. 2C), indicating that ADM mitigates endothelial inflammation, at least in part, through inhibition of NF-κB signaling. Moreover, ADM treatment led to a significant reduction in the soluble ICAM-1, especially at 100 ng/mL (Fig. 2D). Together, these results demonstrate that ADM exerts potent anti-inflammatory effects in endothelial cells exposed to TNF-α, likely via suppression of NF-κB activation.

**Figure 2.**
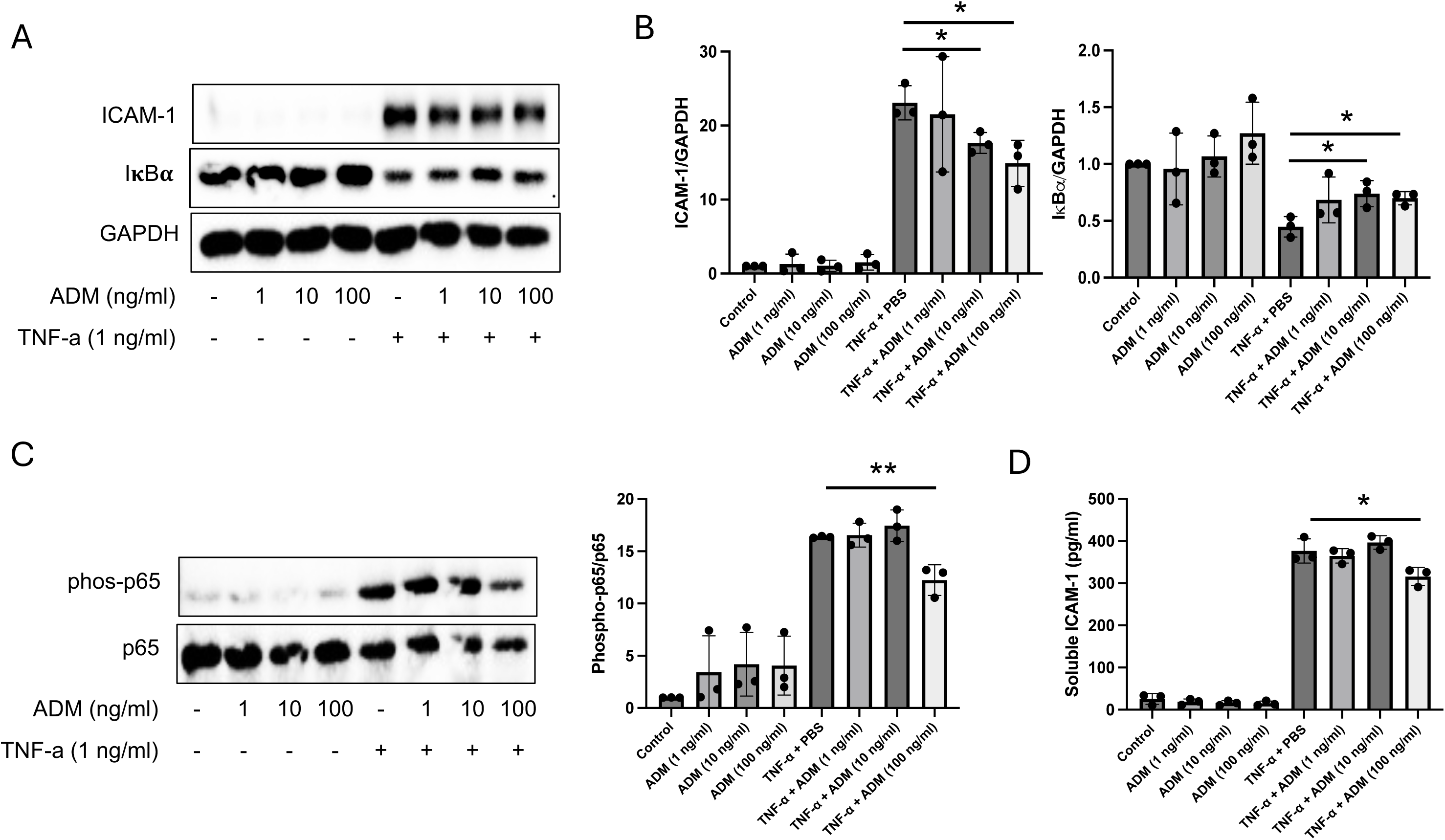
ADM attenuates pro-inflammatory effect of TNF-α. (A and C) HPAECs were treated with TNF-α (1 ng/ml, 30 min) followed by the ADM addition at indicated concentrations for 5.5 hours. The samples were subjected to Western blot (WB) analysis to detect ICAM-1, IκBα, phosphorylated-p65 (S536), total p65, and GAPDH proteins. (B) ICAM-1 and IκBα proteins were normalized to GAPDH, and the values were presented as a bar graph. (C) Phosphorylated-p65 (S536) expression was normalized to total p65 protein levels. (D) The cultured medium from cells treated with 1 ng/ml TNF-α for 30 minutes, followed by ADM for 5.5 hours, was used to measure soluble ICAM-1 expression levels by ELISA assay. The graphs are presented as the means ± standard deviation (SD) from three independent experiments. **\***p<0.05 and **\*\***p<0.01

### 3.3. ADM improves endothelial homeostatic markers under shear stress

Pulmonary endothelial cells are continuously exposed to hemodynamic forces that are essential for maintaining cellular phenotype and physiological functions. To examine the effects of shear stress on endothelial behavior, we applied unidirectional laminar flow using an Ibidi system. After 48 hours of exposure to a shear stress of 15 dynes/cm^2^, HPAECs exhibited increased expression of the mechanotransduction transcription factors *KLF2* and *KLF4*, along with upregulation of canonical endothelial markers *CD31*, *CDH5*, *NOS3*, *KDR*, *TEK*, and *S1PR1* (Figs. 3A and B), suggesting an improvement of endothelial characteristics and homeostasis.

**Figure 3.**
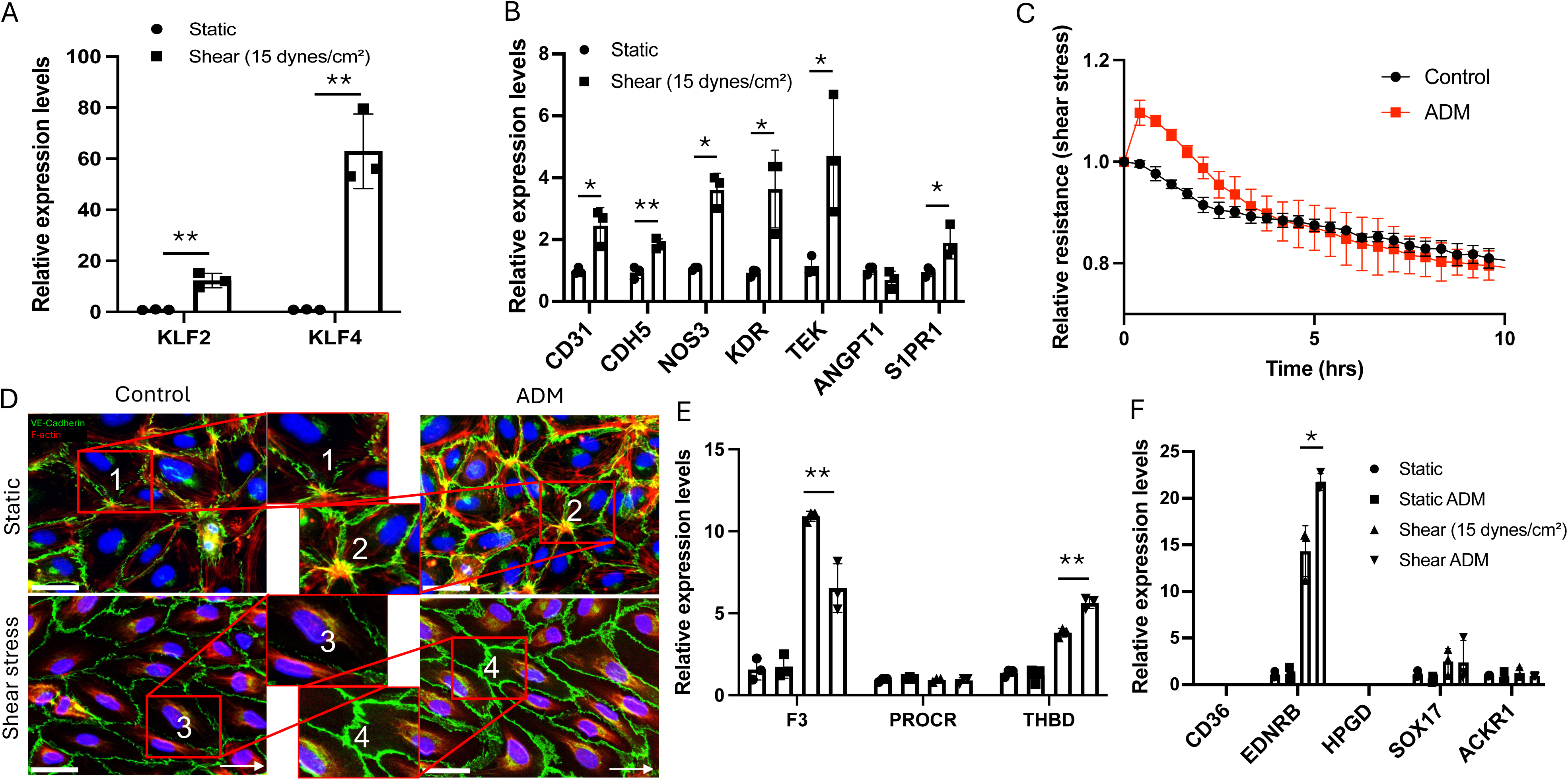
ADM and physiological shear stress synergistically improve endothelial characteristics. (A and B) HPAECs were incubated under unidirectional flow at 15 dynes/cm^2^ for 1 day. The expressions of mechanosensory genes (*KLF2* and *KLF4*) and pan-endothelial cell marker genes (*CD31*, *CDH5*, *NOS3*, *KDR*, *TEK*, *ANGPT1*, and *S1PR1*) were amplificated by real time PCR, normalized to *GAPDH*, and presented as graphs. (C) The cell barrier function was measured by ECIS assay. The graph shows the results from the cells treated with 100 ng/ml ADM after 1 day of shear stress at 15 dyne/cm^2^. (D) The fluorescence images were taken after the cells were treated with 100 ng/ml of ADM for 1 hour under both static and shear stress conditions. For shear stress condition, the cells were incubated for 1day at 15 dynes/cm^2^ before the ADM treatment. (E and F) The cells were incubated with either static or shear stress condition for 1 day, and then 100 ng/ml of ADM was added to the cells after medium change for 30 minutes. The indicated gene expressions were measured by real-time PCR and normalized to GAPDH. The graphs are presented as the means ± standard deviation (SD) from three independent experiments. White scale bars: 50 μm, **\***p<0.05, **\*\***p<0.01, and **\*\*\***p<0.001

To further assess the effect of ADM on endothelial barrier function under shear stress, ADM (100 ng/mL) was added on day 2 of flow culture, and transendothelial electrical resistance was monitored over 10 hours. Consistent with the findings under static culture, ADM markedly enhanced endothelial cell barrier function under shear conditions (Fig. 3C). The reorganization of actin cytoskeleton and adherens and tight junction proteins is closely linked to cell-cell junction regulation^13^. Immunofluorescence images showed that cells under static culture were randomly oriented, with F-actin filaments located at both the cell borders and the perinuclear region, whereas under shear conditions, the cell morphology was changed, and F-actin filaments were reorganized in the direction of flow and significantly diminished at the cell borders (Fig. 3D), suggesting the significant impact of shear stress on actin cytoskeleton reorganization. ADM treatment further reorganized VE-cadherin along the cell borders, promoting a more linear and thicker arrangement that led to the structural reinforcement and stabilization of cell-cell junctions in both shear and static conditions (Fig. 3D). These findings suggest that ADM reinforces endothelial barrier function under shear stress, potentially through cytoskeletal remodeling and stabilization of adherens junctions.

We further assessed the impact of ADM on the expression of endothelial homeostasis, inflammation, and coagulation-related gene expression under shear conditions. We found that ADM had modest or no impact on the improved level of endothelial markers such as *TEK*, *ANGPT1*, and *S1PR1* (Supplemental Fig. 1A) under shear conditions. Shear stress at 15 dynes/cm^2^ alone modestly changed the level of pro-inflammatory markers including *IL1B*, *IL6*, and *ICAM1* (Supplemental Fig. 1B), but strongly affected coagulation-related genes. Shear stress increased coagulation factor III (*F3*; a pro-coagulant factor) and thrombomodulin (*THBD*; an anti-coagulant enzyme) (Fig. 3E). ADM treatment shifted this balance toward an anti-coagulative state by decreasing *F3* and increasing *THBD* (Fig. 3F), suggesting that while shear stress regulates pro- and anti-coagulant balance, ADM promotes an anti-coagulation phenotype.

It is well known that endothelial cells (ECs) tend to lose their native, tissue-specific phenotypes and adopt more generic characteristics when cultured under static in vitro conditions using standard commercial media ^8,14^. We therefore evaluated the impact of ADM and shear stress on endothelial phenotypic markers ^8,15^. Specifically, we assessed the canonical markers for EC subpopulation in the lung vasculature, including *EDNRB* and *HPGD* for aCap, *CD36* for gCap, *SOX17* for arterial, and *ACKR1* for venous endothelial cells. Shear stress alone significantly increased the expression of *EDNRB*, while modestly increased on *SOX17* and having no impact on *ACKR1*. Notably, ADM treatment for 24 hours under shear conditions synergistically enhanced *EDNRB* expression (Fig. 3F), with no impact on other genes. These results suggest that shear stress alone can partially restore endothelial subtype-specific gene expression, while ADM further amplifies this effect, particularly in aCap-associated markers. Collectively, these findings indicate that shear stress promotes endothelial homeostasis and specification, and that ADM further enhances this process by favoring an anticoagulant endothelial state and upregulating EDNRB.

### 3.4. ADM regulates endothelial cell functions through an RAMP2-dependent mechanism

To further determine the signaling regulation of ADM on endothelial function, we first examined the expression levels of ADM receptor under both static and shear conditions.

Interestingly, receptor activity-modifying protein (RAMP2) and calcitonin receptor-like receptor (CALCRL), co-receptors for ADM, were significantly increased under flow, with RAMP2 increasing the most (Fig. 4A). ADM treatment did not alter the expression of either RAMP2 or CALCRL. To further determine the impact of RAMP2 on ADM-induced endothelial maturation, RAMP2 was knocked down with siRNA, followed by ADM treatment. We found that ADM’s impact on endothelial cell barrier protection was almost completely abolished under both static and shear stress conditions (Figs. 4B and C). Furthermore, the ADM-induced VE-cadherin reorganization along the cell borders was also eliminated after RAMP2 depletion (Fig. 4D). The regulation of *EDNRB*, *F3*, and *THBD* gene expression by ADM under shear stress was reduced in RAMP2 knockdown cells (Fig. 4E). Thus, these data demonstrate that ADM-induced effects are dependent on RAMP2, in which the expression level is enhanced by shear stress. These data further suggest that basal RAMP2 levels under static conditions are insufficient to support robust ADM signaling.

**Figure 4.**
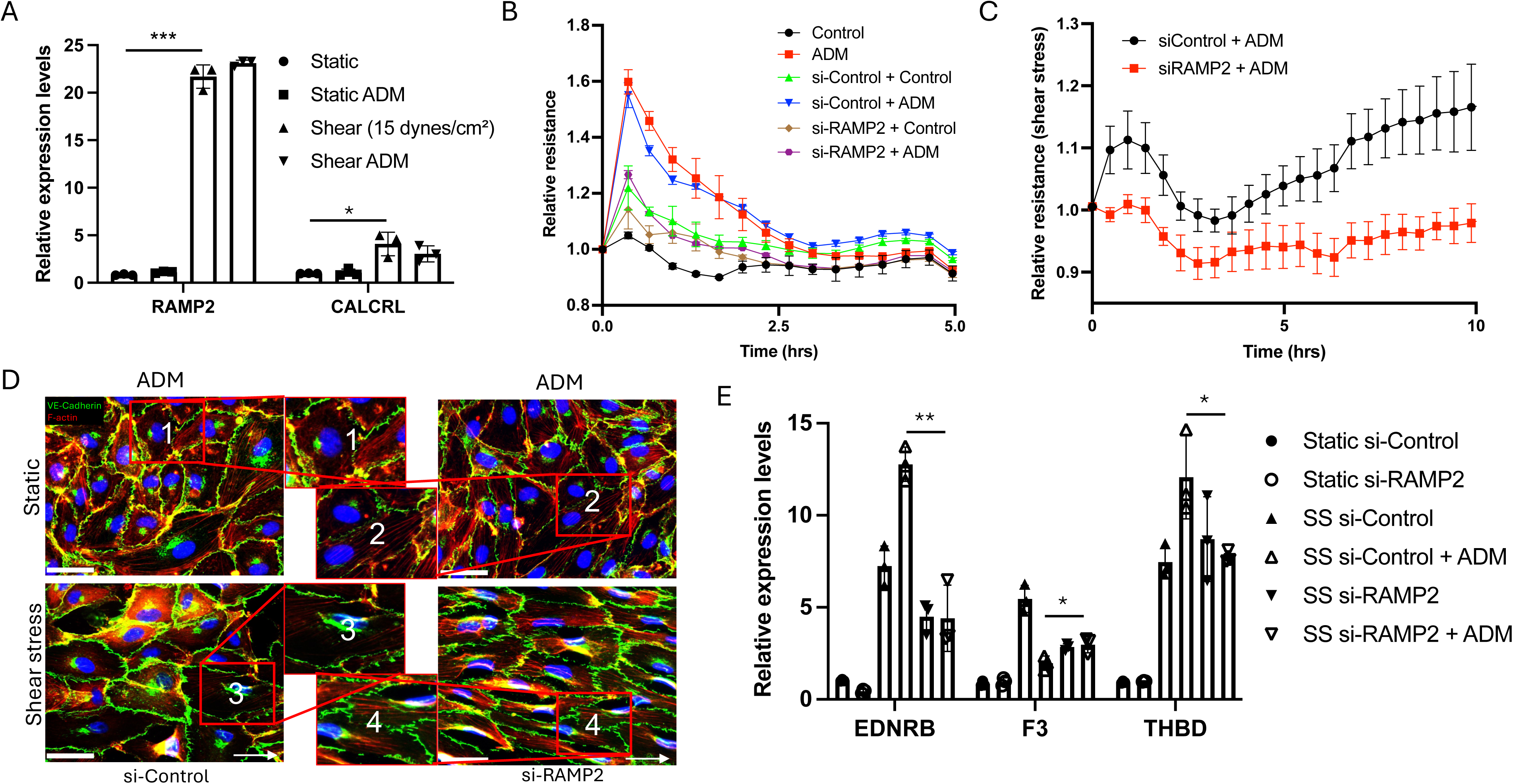
Knockdown of RAMP2 attenuates ADM-mediated enhancement of barrier function and normalization of pulmonary EC phenotype. (A) mRNA expression of ADM target receptors, including RAMP2 and CALCRL, was measured under static and shear stress conditions in the presence of ADM treatment. To knock down of RAMP2 gene expression, control or RAMP2 siRNA (10 nM) were transfected into HPAECs. The ability of ADM (100 ng/ml) to enhance endothelial cell barrier function was measured under both (B) static and (C) shear stress (15 dyne/cm^2^) conditions. (D) The fluorescence images captured from the cells treated with ADM (100 ng/ml) for 1 hour, under both static and shear stress (15 dyne/cm^2^) conditions after transfection of control or RAMP2 siRNA. (E) The genes EDNRB, F3, and THBD, which are regulated by ADM treatment under shear stress, were assessed in the RAMP2 knockdown condition. The graphs are presented as the means ± standard deviation (SD) from three independent experiments. **\***p<0.05 and **\*\***p<0.01

## 4. Discussion

In this study, we compared the impact of different ligands that were previously identified in human lung vascular niche on endothelial functions and phenotypes. We determined that ADM exerts a superior impact on barrier enhancement in human lung endothelial cells compared with other ligands tested in our cell culture model (Fig. 1). We further showed that ADM treatment reduced inflammatory responses under the challenge with TNF-α. Under a shear stress of 15 dynes/cm^2^, ADM preserved barrier improvement and exerted an anticoagulation effect while reinforcing native endothelial characteristics. Therefore, paracrine signals identified from native vascular niche, such as ADM, might contribute to homeostasis in vitro vascular model system, and this favorable impact may not be limited to just vascular engineering alone.

Leveraging scRNA-seq, we previously identified a group of ligand-receptor pairs that are enriched in the vascular niche in the healthy adult human lungs ^8^. However, the impact of these factors on endothelial cell behaviors has not been studied. In the current study, we applied two different assays (ECIS and XperT) to compare the effects of different ligands on endothelial cell barrier function. Our data indicated that Ang-1 and ADM significantly decreased endothelial permeability in both assays, consistent with previous literature ^16,17^. Interestingly, we observed that ADM had a significantly greater effect on improving the vascular barrier as compared to other molecules, which prompts us to focus on ADM in further studies.

Physiological shear can stimulate endothelial homeostasis and reduce inflammatory and coagulation signals ^18–20^. Under hemodynamic forces, especially laminar flow, quiescence and anti-inflammatory phenotypes are induced through multiple mechanotransduction pathways, including KLF2/KLF4, Nrf2, AMPK, and SIRT1 cascades ^21–25^. After two days of dynamic flow culture using an ibidi microfluidic system, we observed significant upregulation of *KLF2* and *KLF4*. Pan-endothelial markers including *CD31*, *CDH5*, *NOS3*, *KDR*, *TEK*, and *S1PR1* were also markedly increased (Fig. 3B), consistent with improvement of a quiescent endothelial phenotype ^26–28^. Hemodynamic forces could also activate mechanosensitive channel PIEZO1, leading to ADM release from endothelial cells ^29,30^. ADM subsequently activates CALCRL in complex with RAMP2, triggering Gs-cAMP-PKA signaling that both attenuates NF-κB-dependent mechanism ^30^ to reduce inflammation and phosphorylates eNOS at Ser633/635 and S1177 to resolve blood clot formation, thereby maintaining endothelial homeostasis ^29^. These data suggest a potential interplay between shear stress and ADM signaling in promoting endothelial homeostasis.

Pulmonary vascular diseases, including sepsis, pneumonia, and pulmonary hypertension, are characterized by inflammation and activation of the coagulation cascade ^31–33^. Low or absent shear promotes the formation of red blood cell (RBC)- and fibrin-rich(“red”) thrombi, whereas higher-shear arterial condition favor platelet-rich (“white”) thrombi ^31,34,35^. Thrombomodulin is a transmembrane glycoprotein expressed on endothelial cells and serves as a cofactor for thrombin-mediated activation of protein C to APC, which has anticoagulant, anti-inflammatory, and anti-apoptotic effects ^36^. Previous study demonstrates that laminar flow enhanced THBD expression in human brain microvascular endothelial cells ^37^. Consistently, our results show that physiological shear stress increased *THBD* expression, and modulated the coagulation profile, with an increase in *F3* (tissue factor), the initiator of the extrinsic coagulation pathway, while endothelial protein C receptor (*PROCR*) remained unchanged, suggesting that shear stress at 15 dynes/cm^2^ may regulate a balance between pro- and anti-coagulant states. In the presence of shear stress, ADM shifted the endothelial phenotype toward an anticoagulant state as evidenced by increased THBD and decreased F3, whereas no such effect was observed under static conditions (Figs. 3E and 4E). Cyclic AMP signaling has been reported to regulate coagulation and inflammation by downregulating F3 and upregulating THBD across monocytes, cancer, vascular smooth muscles, and endothelial cells, and this signal has been shown to be regulated through ADM-RAMP2 signaling ^38–42^. In line with this, RAMP2 knockdown abrogated ADM-induced modulation of F3 and THBD (Fig. 4E), consistent with engagement of the ADM-CALCRL/RAMP2-cAMP pathway.

There are two major endothelial cell populations within lung microvascular bed, aCap, which are specialized for gas exchange and immune cell recruitment, and gCap, which regulate vasomotor tone and serve as progenitors in the lung microvasculature ^15^. We found that shear stress alone (15 dynes/cm^2^) could markedly increase *EDNRB* expression, a critical marker for aCap cells, and this effect was further enhanced by ADM treatment. Both hemodynamic forces and ADM have been reported as critical cues contributing to vasodilation in the lung vasculature. EDNRB is known to mediate vasodilation upon binding with endothelins (ET-1, ET-2 and ET-3) ^43^. Previous studies have reported that ADM enhances ET-1 release and promotes ET-1-EDNRB interactions in rat lung endothelial cells ^44^. Together, these results suggest that ADM might synergistically enhance vasodilatory responses through an ET-1/EDNRB-dependent mechanism under shear stress.

There are several limitations associated with this project: 1) We primarily compared the effects of different ligands on endothelial barrier function and subsequently focused on ADM for detailed analysis. This prioritization was based on ADM’s strong and reproducible effects on barrier integrity, which suggested a robust biological role. As a result, other ligands were not comprehensively evaluated for their anti-inflammatory, anticoagulant, or native endothelial phenotype-related effects. Future studies will systematically assess these additional functional dimensions. 2) Another limitation is that our shear stress experiments were conducted at a single physiological condition (15 dynes/cm^2^), chosen based on prior literature describing typical physiological shear levels. While this approach allowed us to directly compare results with published findings, it limited our ability to examine potential dose- or pattern-dependent effects of shear stress. Subsequent studies will explore a broader range of shear conditions to better capture the mechanobiological spectrum of endothelial responses.

Together, our data support that ADM responsiveness is shear-dependent, in which shear enables robust effects, whereas static condition does not, underscoring shear’s role as an integrator of mechanotransduction and paracrine inputs. Whether ADM’s shear-dependent effects persist across different shear stress magnitudes and endothelial subtypes remains to be determined, a question with direct implications for how we model endothelial homeostasis and test soluble cues in vitro.

