## Supplemental Fig 1 for "Adrenomedullin-RAMP2 enhances endothelial cell homeostasis synergically with shear stress"

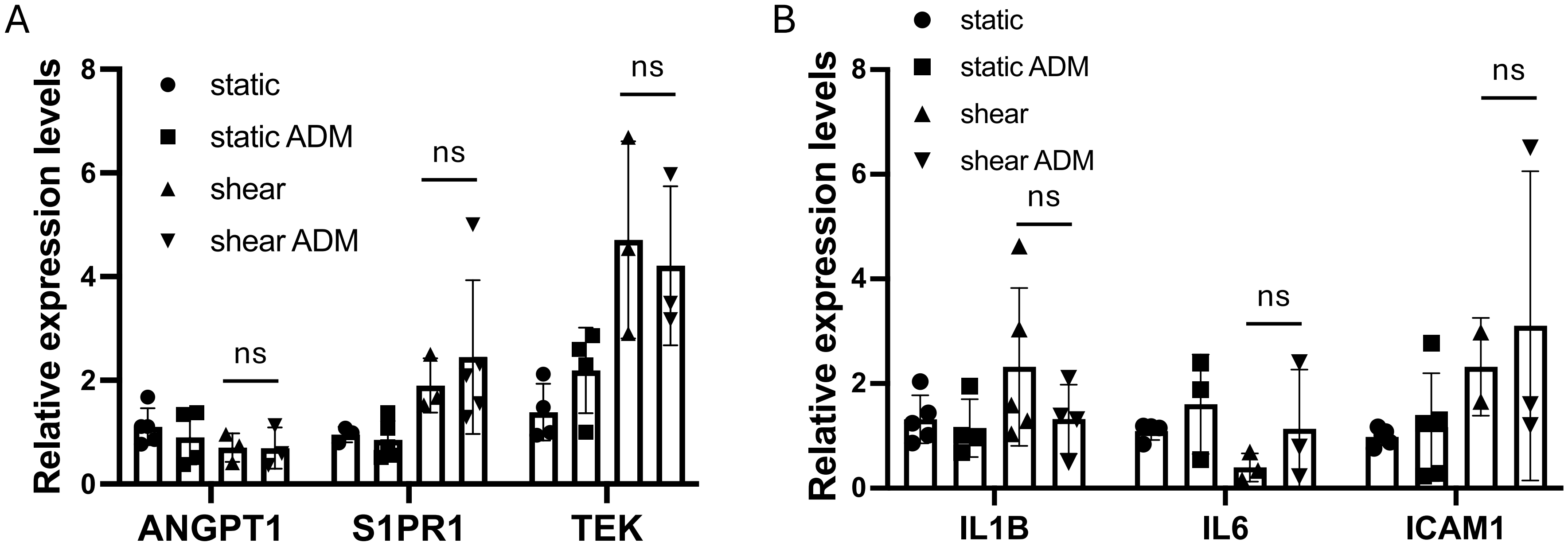


**Supplemental Figure 1. ADM and endothelial characteristics under shear stress.** (A and B) The cells were incubated under either static or shear stress condition for 1 day, followed by the treatment of ADM. The samples were collected after 24 hours of treatment. The expression level of *ANGPT1*, *S1PR1*, *TEK*, *IL1B*, *IL6*, and *ICAM1* were evaluated. The graphs are presented as the means ± standard deviation (SD) from three independent experiments. ns indicates non-significant.
